# CXCR4 coordinates adhesion, migration, and development of human NK cells

**DOI:** 10.64898/2026.03.12.711152

**Authors:** Shira E. Eisman, Francesca Grossberg, Batya S. Koenigsberg, David H. McDermott, Frédérique van den Haak, Luis A. Pedroza, Everardo Hegewisch-Solloa, Philip M. Murphy, Emily M. Mace

## Abstract

Natural killer (NK) cells undergo stepwise differentiation from multipotent progenitors within secondary lymphoid tissues. Despite the central importance of the tissue microenvironment in their development, little is known about cell-cell interactions that regulate human NK cell trafficking and maturation. Here, we identify the chemokine receptor CXCR4 and its ligand CXCL12 as regulators of stromal-NK cell interactions required for NK cell maturation. We demonstrate that CXCR4 is expressed throughout human NK cell development in peripheral blood and tonsil, and CXCL12 is enriched in stromal niches containing developing NK cells. Pharmacologic blockade or genetic disruption of CXCR4 resulted in diminished adhesion to integrin ligands and high-resolution imaging demonstrated crosstalk between CXCR4 and integrins, providing a mechanistic basis for chemokine-dependent modulation of adhesion. Further, CXCR4 blockade resulted in altered contact-dependent motility on stromal cells and integrin ligands, with decreased stable stromal engagement and increased cell speed. Consistent with a requirement for these interactions, treatment with the CXCR4 antagonist plerixafor (AMD3100) impaired NK cell generation from CD34^+^ precursors. Analysis of NK cells from WHIM syndrome patients with CXCR4 gain-of-function mutations treated with plerixafor revealed similar defects in migration and adhesion, supporting the in-vivo relevance CXCR4-dependent regulation of NK cell adhesion and motility.

## Introduction

Natural Killer (NK) cells are innate lymphocytes that contribute to early protection against viral infection and malignancy by exerting effector functions without the need for antigen presentation (1). Classically defined as CD3^−^-CD56^+^ granular lymphocytes, NK cells originate from hematopoietic stem cells (HSCs) in the bone marrow (BM) and undergo stepwise maturation that can be defined by distinct surface markers and stage-specific functions (2, 3).

While recent studies show that early NK precursors leave the BM and mature in secondary lymphoid tissues such as tonsil (4–7), the cellular interactions that guide human NK development remain incompletely understood. Fetal liver-derived stromal cell lines support NK cell differentiation in-vitro from CD34^+^ precursors (8–10), and direct stromal contact effectively promotes human NK maturation (11). While the molecular signals governing these interactions remain unclear, these observations point to a key role for stromal microenvironments that coordinate NK polarization, adhesion, and developmental progression. Previous studies of NK cells isolated from human peripheral blood show that they exhibit non-directed cell motility when plated on developmentally supportive stroma (12). NK cell subsets display distinct migratory behaviors on stromal cells, suggesting an intricate link between migration and development. A specialized NK-stroma contact termed the developmental synapse (DS) has been described and is marked by actin polarization, CD56 accumulation, and signaling activity (12). Together, these findings support a model in which regulated NK-stromal interactions contribute to human NK maturation, but the upstream cues that stabilize or modulate these contacts remain unknown.

Chemokine-receptor interactions likely stabilize these contacts. CXCR4 and its ligand CXCL12 regulate hematopoietic cell retention, homing, and tissue organization. Beyond their well-described role in chemotaxis (13, 14), CXCR4-CXCL12 can also promote synapse formation and polarization in developing immune cells (15). While CXCR4 has not been studied extensively in human NK cells, murine studies demonstrate its importance, as hematopoietic lineage specific CXCR4 conditionally deficient mice display decreased NK precursors in the BM and fewer mature NK cells in peripheral tissues (16). In addition, NK cells in mice localize near CXCL12-abundant reticular (CAR) cells that provide IL-15, an essential cytokine for NK development (17). These observations raise the possibility that CXCR4-CXCL12 actively shapes NK cell maturation through mechanisms beyond chemotaxis.

Aberrant leukocyte trafficking is a defining feature of primary immunodeficiencies that include NK cell abnormalities, including Wiskott-Aldrich syndrome and GATA2 deficiency (18, 19). Among disorders directly linked to dysregulated chemokine signaling, WHIM syndrome, a primary immunodeficiency defined by Warts, Hypogammaglobulinemia, Infections, and Myelokathexis, results from gain-of-function mutations in CXCR4 that truncate its C-terminal tail, impairing receptor internalization and causing sustained signaling (20–22). This hyperactive CXCR4 signaling promotes leukocyte retention in the BM, pan-leukopenia, and recurrent infections (23–25). The CXCR4 antagonist plerixafor (AMD3100), which is used clinically for hematopoietic stem-cell mobilization, has incompletely understood effects on NK cell development and function (26, 27).

Here, we show that CXCR4 regulates human NK cell development by stabilizing integrin-dependent interactions with stromal cells. We demonstrate that CXCR4-CXCL12 interactions promote adhesion, polarization, and developmental progression of human NK cells, thus linking NK cell engagement with stromal cells to human NK cell maturation.

## Results

### CXCR4 is expressed on immature and mature human NK cells

To investigate whether CXCR4-CXCL12 signaling might play a role in human NK cell development, we first assessed CXCR4 expression across NK cell developmental stages by examining CXCR4 transcript levels in circulating lymphoid progenitors and NK cell subsets using data from the Human Immune Health Atlas (28, 29). CXCR4 was detected in the ILC (innate lymphoid cell) progenitors detectable in PBMC (peripheral blood mononuclar cells) in this dataset, CD56^bright^ NK cells, CD56^dim^ NK cells, and proliferating NK cell subsets, with the highest mean expression in CD56^dim^ NK cells (Fig.1A).

We next assessed CXCR4 surface expression by flow cytometry on NK cell subsets and NK progenitors from tonsil, a site of human NK cell maturation (3) (Fig. 1B; gating strategy in Supp. Fig. 1A). As predicted by its association with early hematopoietic differentiation (30, 31), we found significantly higher CXCR4 expression in CD34^+^ hematopoietic stem and progenitor cells (HSPCs) from tonsil when compared to mature CD16^−^ NK cells (Fig. 1C, D). Additionally, CD16^+^ NK cells from tonsil show a significant increase in CXCR4 mean fluorescence intensity (MFI) relative to CD16^−^ NK cells (Fig. 1C, D), confirming that the higher expression of CXCR4 transcript in CD56^dim^ NK cells (Fig. 1A) was also associated with higher protein expression.

**Fig. 1.**
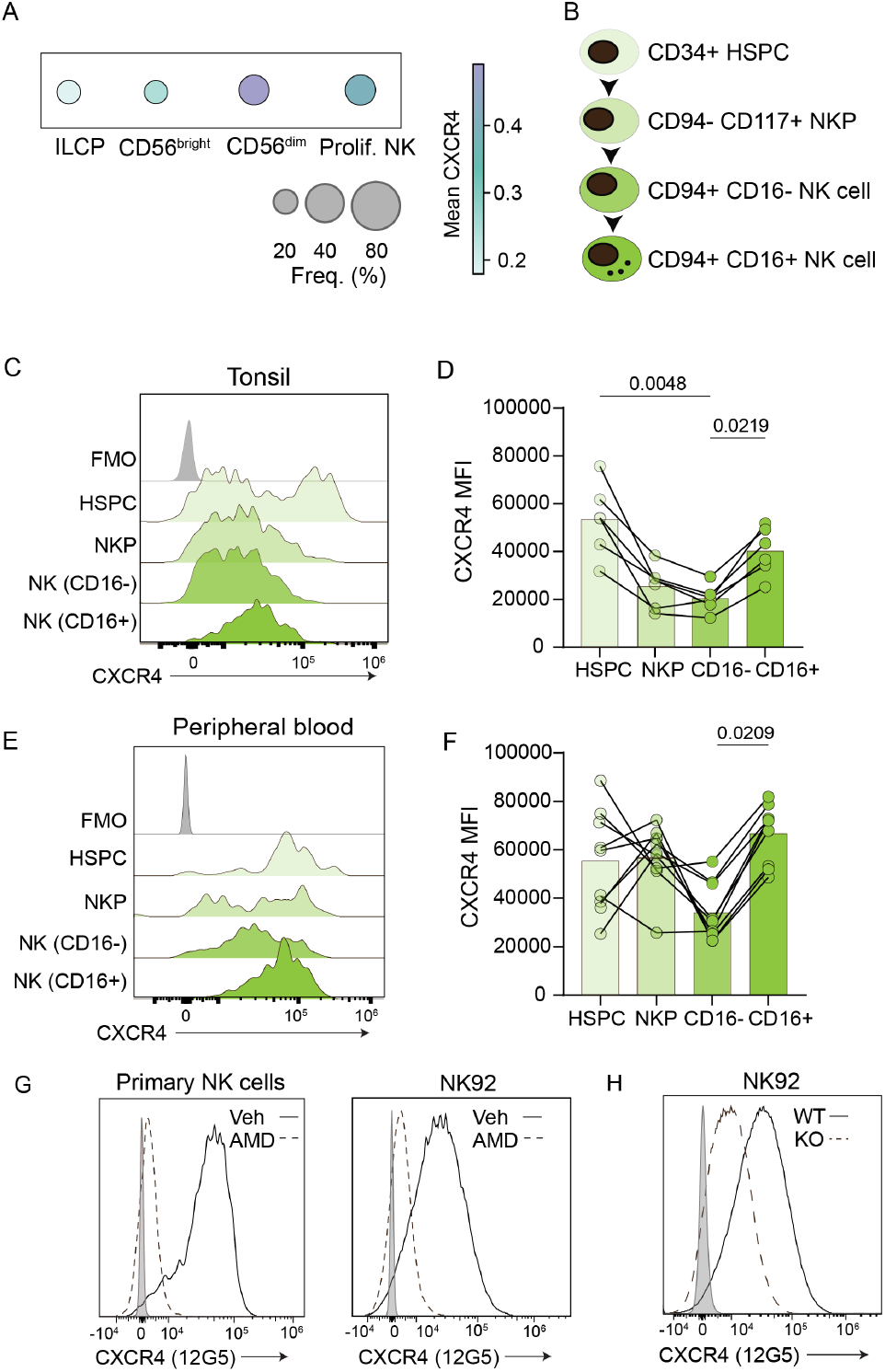
CXCR4 is expressed throughout human NK cell development. A) CXCR4 transcript levels in ILC-like innate lymphoid cell progenitors (ILCPs) and NK subsets from the Human Immune Health Atlas (29) The earliest population represents the ILC-labeled innate lymphoid population. Dot size indicates frequency of CXCR4^+^ cells; color scale represents mean expression. B) Schematic of NK cell developmental progression: CD34^+^ hematopoietic stem and progenitor cells (HSPCs), CD117^+^CD94^−^ NK precursors (NKP), CD94^+^CD16^−^ NK cells, and CD94^+^CD16^+^ NK cells. C) Representative histograms showing mean fluorescence intensity (MFI) of extracellular CXCR4 in NK developmental intermediates from tonsil. Gray histogram = Fluorescence minus one (FMO) control. D) Pooled data from 6 healthy donors corresponding to flow cytometry data in (C). Each dot represents an individual donor (n=6), and lines connect populations derived from the same donor. Horizontal bars indicate the median. Statistical significance was determined using a Friedman test with Dunn’s multiple-comparisons correction (paired by donor). Only p<0.05 are displayed. E) Representative histograms showing extracellular CXCR4 MFI in NK developmental intermediates from peripheral blood. F) Pooled data from 9 healthy donors corresponding to flow cytometry data in (E). Each dot represents an individual donor (n=9), and lines connect populations derived from the same donor. Horizontal bars indicate the median. Statistical significance was determined using a Friedman test with Dunn’s multiple-comparisons correction (paired by donor). Only p<0.05 are displayed. G) Representative histograms of cell surface CXCR4 MFI in primary NK cells (left) and NK92 cell lines (right) after 1hr treatment with Vehicle (Water; solid) as a control or 10 *µ*M AMD3100 (dashed). Representative of >10 biological replicates. H) Validation of CXCR4 knockout (KO) in NK92 cell line. Representative histogram of CXCR4 staining in wild-type (WT; solid) and CRISPR-Cas9 CXCR4-KO (dashed) NK92 cells. Representative of 4 technical replicates.

We then performed the same analysis using peripheral blood. Although CD34^+^ HSPCs are rare in peripheral blood, a similar trend towards higher expression in CD34^+^ HSPCs was observed relative to CD16^−^ NK cells (Fig.1E, F). Circulating NK/ILC precursors (Lin^−^CD117^+^CD94^−^) in peripheral blood (32) showed comparable CXCR4 expression to HSPCs. Similarly to tonsil, CD16^+^ NK cells displayed higher CXCR4 MFI than CD16^−^ NK cell subsets (Fig 1E, F). Together, these flow cytometry data demonstrated that CXCR4 is expressed at all stages of human NK cell development in peripheral blood and tonsil, with differences in expression associated with changes in NK cell maturation and location.

To establish experimental models for CXCR4 perturbation, we used pharmacologic blockade with a CXCR4 antagonist (AMD3100) on primary NK cells and generated a CRISPR-Cas9-mediated CXCR4 knockout (KO) in the NK92 NK cell line. As expected, AMD3100 treatment of primary NK cells and NK92 cells reduced CXCR4 detection by flow cytometry due to competitive binding with the CXCR4-specific antibody clone 12G5 (Fig. 1G) (33, 34). Targeted knockout of CXCR4 using CRISPR-Cas9 results in a single population with 50% decreased CXCR4 protein expression by flow cytometry (Fig. 1H). Neither pharmacological blockade nor genetic deletion affected NK-cell viability or proliferation (Supp. Fig.1B). In transwell assays, both AMD3100 and CXCR4 knockout significantly impaired directed migration toward CXCL12, confirming that both approaches impacted the canonical function of CXCR4 in chemotaxis (Supp. Fig. 1C). Decreased chemotaxis in the CXCR4-KO cells was moderate but could be further reduced in the presence of AMD3100 (Supp. Fig 1C). In contrast, NK cell cytotoxic function in a standard 4-hour killing assay against K562 target cells was only modestly reduced by CXCR4 knockout (Supp. Fig. 1D) or AMD3100 (Supp. Fig. 1E). Together, these results demonstrate that CXCR4 is expressed throughout NK cell development and in NK cell lines, and that it is not required for NK cell survival or cytotoxic function.

### CXCR4 ligand CXCL12 is found in NK cell developmental niches

CD34^+^ progenitors in peripheral blood that give rise to NK cells are thought to enter the tonsil through high endothelial venules (HEVs) within the interfollicular domain, then migrate to the parafollicular domain where they undergo further maturation (3). As CXCL12 is the primary ligand for CXCR4, we sought to define its spatial localization in tonsil using cyclic immunofluorescence (CycIF) on formalin-fixed, paraffin embedded sections from routine pediatric tonsillectomies (35).

Observing CXCL12 expression in CycIF images, we found that CXCL12 protein was distributed throughout the tonsil but enriched in inter- and parafollicular stromal regions surrounding follicles and in HEVs (Fig. 2A). Quantification of multiple regions of interest from samples from multiple donors confirmed higher CXCL12 intensity in inter- and parafollicular regions relative to B cell follicles (Fig. 2B). Using our multiplexed images, we defined CD56^+^CD3^−^ NK cells and used Ki67 to visualize proliferating cells (Fig. 2C, additional channels shown in Supp. Fig. 2). High magnification images showed that NK cells were often adjacent to CXCL12^+^ stromal cells; and many, but not all, of these were Ki67+ (Fig. 2Ci-iii). Further analysis revealed that CD34^+^ progenitors were also found adjacent to CXCL12^+^ stroma (Supp. Fig. 2Aiii).

**Fig. 2.**
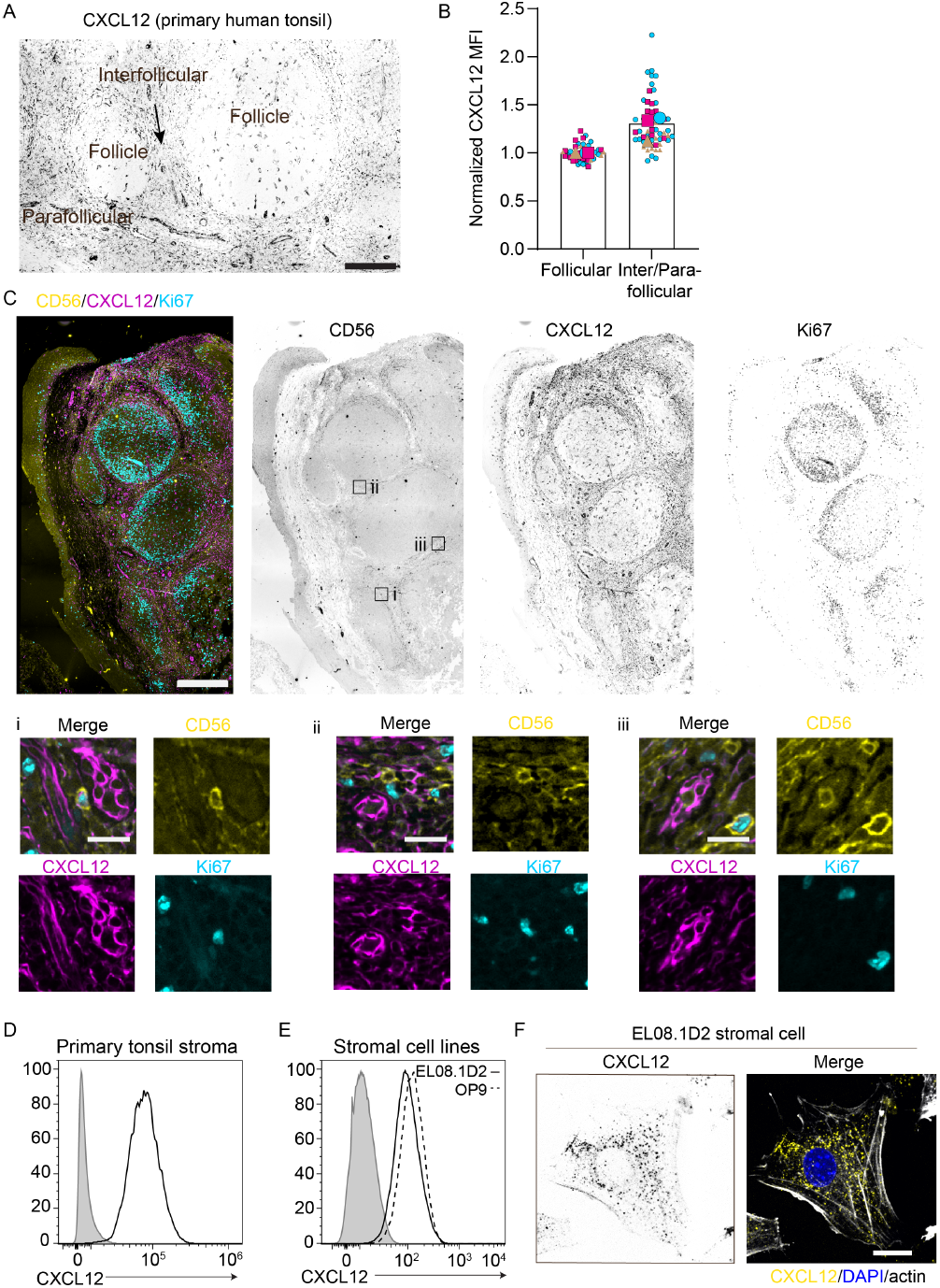
CXCL12 is found in sites that support human NK cell development. To define the localization of NK cell subsets relative to CXCL12-expressing cells, we performed cyclic immunofluorescence on pediatric human tonsil. A) Representative immunofluorescence staining of CXCL12 in primary human tonsil tissue with follicles and interfollicular and parafollicular regions annotated. B) Quantification of normalized CXCL12 mean fluorescence intensity (MFI) comparing follicular versus para- and interfollicular regions; each point represents a region of interest from 3 donors, 8-12 regions per donor. Large symbols represent mean of values from individual donors. C) Cyclic immunofluorescence staining of pediatric tonsil showing CD56^+^ NK cells (yellow), CXCL12 (magenta), and Ki67 (cyan). Grayscale images show individual channels CD56, CXCL12, and Ki67. Scale bar = 100 µm. Boxes denote regions i-iii. Insets show higher-magnification views of NK cell-rich para- and interfollicular stromal regions with CD56^+^ NK cells adjacent to CXCL12^+^ stroma. D) Flow cytometric analysis of primary tonsil stromal cells stained for intracellular CXCL12 (black) compared with fluorescence minus one control (gray). E) CXCL12 expression by stromal cell lines. Flow cytometry comparing intracellular CXCL12 expression on EL08.1D2 (solid line) versus OP9 (dashed line) stromal cells with fluorescence minus one control (gray). F) Confocal imaging of EL08.1D2 stromal cells showing CXCL12 (yellow), DAPI (blue), and actin (white). Scale bar = 5µm.

To further demonstrate that tonsil stromal cells are a source of CXCL12, we isolated CD45-primary tonsil stromal cells from routine tonsillectomies. Flow cytometry demonstrated that primary tonsil stroma uniformly express CXCL12 that was detected following fixation and permeabilization (Fig. 2D). We also found that OP9 and EL08.1D2 cell lines, which support NK cell development, similarly express CXCL12 (Fig. 2E). We confirmed this finding with confocal microscopy of EL08.1D2, which showed intracellular and cell surface expression of CXCL12 (Fig. 2F). These results are consistent with CXCL12 being produced intracellularly for secretion and subsequent binding to the extracellular matrix (36, 37). Together, these data demonstrate that CXCL12 is produced by stromal cells in the inter- and parafollicular zones, forming chemokine-rich microenvironments that likely guide NK progenitors and other CXCR4^+^ cells in the tonsil.

### CXCR4 promotes integrin-mediated interactions between NK cells and stroma

Since NK cell development depends on direct contact with stromal cells and NK cells undergo surveillance-like migration on stromal monolayers in-vitro (12, 38), we asked whether CXCR4-CXCL12 contributes to these migratory behaviors. CD56^+^ NK cells and CD34^+^ progenitors from cord blood were pre-treated with vehicle (water) or AMD3100 and plated on EL08.1D2 stromal monolayers. To assess these interactions in the context of spontaneous cell migration, cells were imaged every 2 minutes for 2-6 hours by live cell confocal microscopy (Fig. 3A). We used Cellpose (39) to segment the cells, btrack (40) to track the cells, and Cell Plasticity Analysis Tool (cellPLATO) (41) to quantify and plot cell motility and shape parameters (Fig. 3B). This analysis focuses on spontaneous contact-mediated motility, in contrast to the directed chemotactic migration measured in transwell assays (Supp. Fig. 1C).

**Fig. 3.**
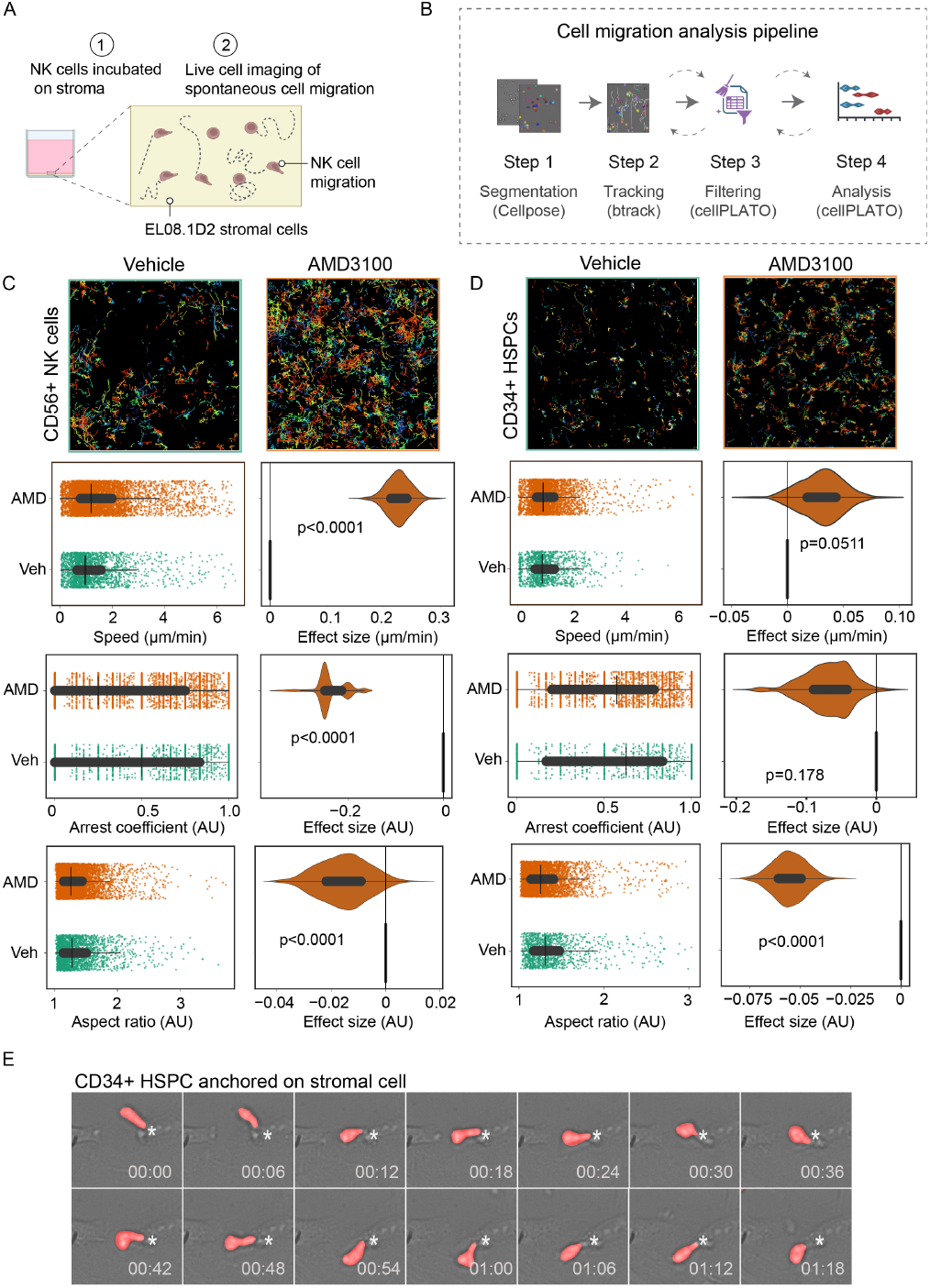
CXCR4-CXCL12 interaction promotes integrin-mediated arrest on stromal cells. Primary CD56^+^ NK cells and CD34^+^ progenitors were plated on EL08.1D2 stromal monolayers or integrin ligand surfaces and imaged by live-cell confocal microscopy every 2 minutes for 2-6 hours to quantify spontaneous, contact-mediated migration. A) Schematic of NK cell migration assay. B) Overview of the cellPLATO migration analysis pipeline. Time-lapse images were segmented using Cellpose, tracked using btrack, and processed through the cellPLATO workflow for track quality control and extraction of motility and morphology features. C) Migration of primary NK cells on stroma with or without AMD3100. Representative segmentation masks and cell trajectories from Vehicle and AMD3100 treated NK cells are shown (top). Color gradient signifies time. Scale bar = 100 *µ*m. Plots of differences show mean speed, arrest coefficient, and aspect ratio. Each point represents a single cell. Effect size calculated by bootstrap resampling. D) Migration of CD34^+^ progenitor cells on stroma with or without AMD3100. Representative segmented images and cell tracks of migrating CD34^+^ cells are shown (top). Color gradient signifies time. Scale bar=100 µm. Migration and shape parameters for CD34^+^ cells with or without AMD3100, including mean speed, arrest coefficient, and aspect ratio. E) Time-lapse sequence of CD34^+^ HSPC with segmentation shown in red interacting with a stromal cell. Asterisk shows the point of contact. Time is shown as hh:mm. Movie shown in Supplemental Movie 2.

We first analyzed the migration of primary NK cells (Fig. 3C). Individual cell trajectories over time were visibly longer and less confined in AMD3100-treated cultures (Fig. 3C, Supp. Movies 1 and 2), suggesting changes in migratory behavior resulting from CXCR4 inhibition. We quantified cell migration parameters, including cell speed and arrest coefficient (the frequency of time a cell is immotile as defined by moving under 1 µm/min). To visualize these differences, we used plots of differences (42) to display the effect size distribution of >8000 cells between conditions. In the vehicle-treated condition, NK cells exhibited characteristic non-directed migration on stromal monolayers, with heterogeneous motility patterns and intermittent arrest (Supp. Movie 1). Overall, AMD3100 treatment increased motility and reduced arrest on stroma (Fig. 3C: Supp. Movie 2). Specifically, AMD3100 treated cells had significantly higher average speeds (1.564 µm/min vs 1.265 µm/min for vehicle) with a reduction in arrest coefficient (0.374 vs 0.483 for vehicle). These data reflect the AMD3100-treated NK cells becoming more motile and less adherent to the stromal surface. Additionally, shape analysis demonstrated rounder, less elongated cells under AMD3100 treatment (aspect ratio of 1.339 vs 1.376 for vehicle). Similar trends were observed in additional biological replicates (Supp. Fig. 3A).

Next, given the expression of CXCR4 on CD34^+^ HSPCs (Fig. 1), we asked whether AMD3100 treatment similarly affects CD34^+^ progenitor cell migration and engagement with stroma (Fig. 3D; Supp. Movies 3 and 4). Although these cells were overall less motile than mature NK cells (12, 43), AMD3100 similarly increased cell speed and modestly reduced arrest coefficient, with rounder and less polarized cells. These data indicate that CXCR4 contributes to progenitor-stroma interactions as well. Consistent with these population-level differences, time-lapse imaging showed cells in the CD34^+^ vehicle-treated condition that remained stably anchored to individual stromal cells for >1 hour, with ongoing protrusive activity but little translocation (Fig. 3E; Supp. Movie 5), a phenotype that has been previously described for HSPCs in contact with bone marrow stroma (15) and was markedly reduced upon AMD3100 treatment (Supp. Movie 6).

To confirm that these observations were not unique to primary cells or pharmacologic blocking of CXCR4, we next analyzed migration of CXCR4 wild type and CXCR4-knockout NK92 cell lines. Like primary NK cells treated with AMD3100, CXCR4 knockout increased cell speed while decreasing arrest coefficient (Supp. Fig. 3B; Supp. Movie 7 and 8).

Since NK cell migration and adhesion are known to be mediated by integrin-ligand interactions (44), we next assessed the migration of primary NK cells with and without AMD3100 on integrin ligands ICAM-1 or VCAM-1 (Supp. Movies 9-12). In both cases, AMD3100 treatment similarly increased motility and reduced arrest, suggesting that CXCR4 signaling directly reinforces integrin-dependent contact with both stromal cells and integrin ligands (Supp. Fig. 3C). Altogether, these findings indicate that CXCR4 contributes to integrin-dependent adhesion and motility.

### CXCR4 promotes integrin activation and NK cell adhesion

To directly test whether CXCR4 modulates NK cell adhesion via integrin activation, we measured adhesion to ICAM-1. Pharmacologic inhibition of CXCR4 significantly reduced the adhesion of primary NK cells to ICAM-1-coated surfaces, and similar reductions were observed in CXCR4-knockout NK92 cells, indicating that CXCR4 activity enhances ICAM-1 engagement (Fig. 4A). Despite the decrease in adhesion, surface expression of CD18 was unchanged (Fig. 4C), indicating that reduced adhesion is not due to changes in integrin surface abundance.

**Fig. 4.**
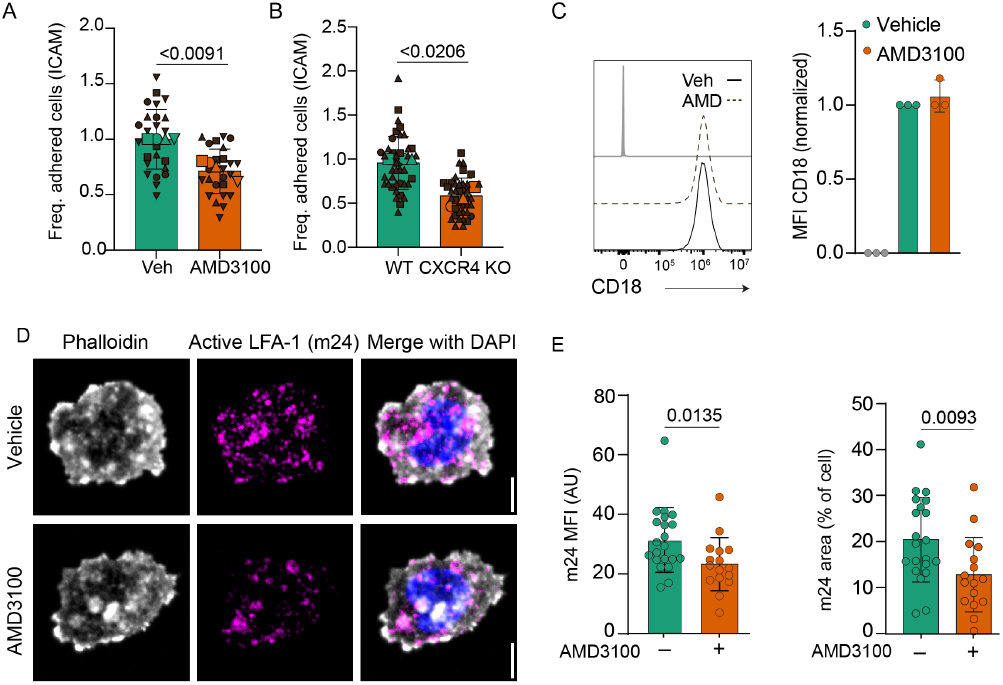
CXCR4 signaling promotes adhesion and integrin activation. A) Cell adhesion assays of primary NK cells on ICAM-1 coated plates following Vehicle or 10 *µ*M AMD3100 treatment. n=4 donors represented by large symbols; mean of each Vehicle treated donor is normalized to 1. 10 regions of interest were imaged per donor and are represented by small symbols. B) Adhesion of NK92 WT or CXCR4-KO cells to ICAM-1 coated plates. n=3 independent experiments; large symbols represent the mean of each technical replicate normalized to 1 for WT condition. 10 regions of interest were imaged per donor and are represented by small symbols. For A and B, adhesion values were normalized to Veh/WT controls within each independent experiment. Each symbol represents a technical replicate. Bars indicate the mean of all replicates. Statistical analysis was performed on experiment means using a two-tailed one-sample t test compared with a theoretical mean of 1. C) Surface CD18 (β2 integrin) expression in primary NK cells after treatment with Vehicle (water) or AMD3100 displayed as representative histograms (left) and normalized MFI (right). Gray=FMO control. n=3 independent experiments (mean±SD). No statistical significance was found using a two-tailed Wilcoxon matched-pairs signed rank test. D) Conformational activation of LFA-1 measured using confocal microscopy. Primary NK cells were treated with 10 *µ*M of AMD3100 or Vehicle and incubated on ICAM-1 (LFA-1 ligand) before fixation and immunostaining with anti-activated LFA-1 (clone m24), phalloidin, and DAPI. Representative confocal images showing phalloidin (actin), m24 (active LFA-1) and merged channels. Scale bar = 5 *µ*M. E) Quantification of m24 MFI and percent area from one representative donor. Each dot represents a single cell from one donor. Horizontal bars indicate mean ± SD. Statistical comparison was performed using a two-tailed Mann–Whitney test across single-cell measurements within this donor.

We therefore examined integrin activation using the conformation-sensitive antibody m24, which recognizes the high affinity state of LFA-1 (45). CXCR4 inhibition with AMD3100 markedly reduced m24 intensity and cluster formation on the NK cell surface following ICAM-1 engagement (Fig. 4D). Quantification of these observations confirmed that AMD3100 significantly decreased the size and intensity of m24 foci (Fig. 4E). Additional donors included in Supp. Fig. 4. Together, these findings demonstrate that CXCR4 signaling promotes β2 integrin activation and ICAM-1-dependent adhesion. This establishes a mechanistic link between CXCR4 and the stabilization of NK cell-stroma interactions, positioning CXCR4 as an upstream regulator of the adhesion-migration balance in human NK cells.

### CXCR4 and CXCL12 are required for effective NK cell differentiation from CD34^+^ cells

Because stromal contact is required for human NK cell maturation, we next asked whether disruption of CXCR4-dependent adhesion affects NK cell developmental progression. To test this, we performed NK cell differentiation from HSPCs in the presence or absence of AMD3100. CD34^+^ cord blood progenitors were isolated by FACS and differentiated on EL08.1D2 stromal monolayers with cytokines (8). Cultures were treated with 10 µM AMD3100 or vehicle (water) at weekly media exchanges (Fig. 5A). At weekly harvests, we quantified total live CD45^+^ cells and NK developmental subsets by flow cytometry.

**Fig. 5.**
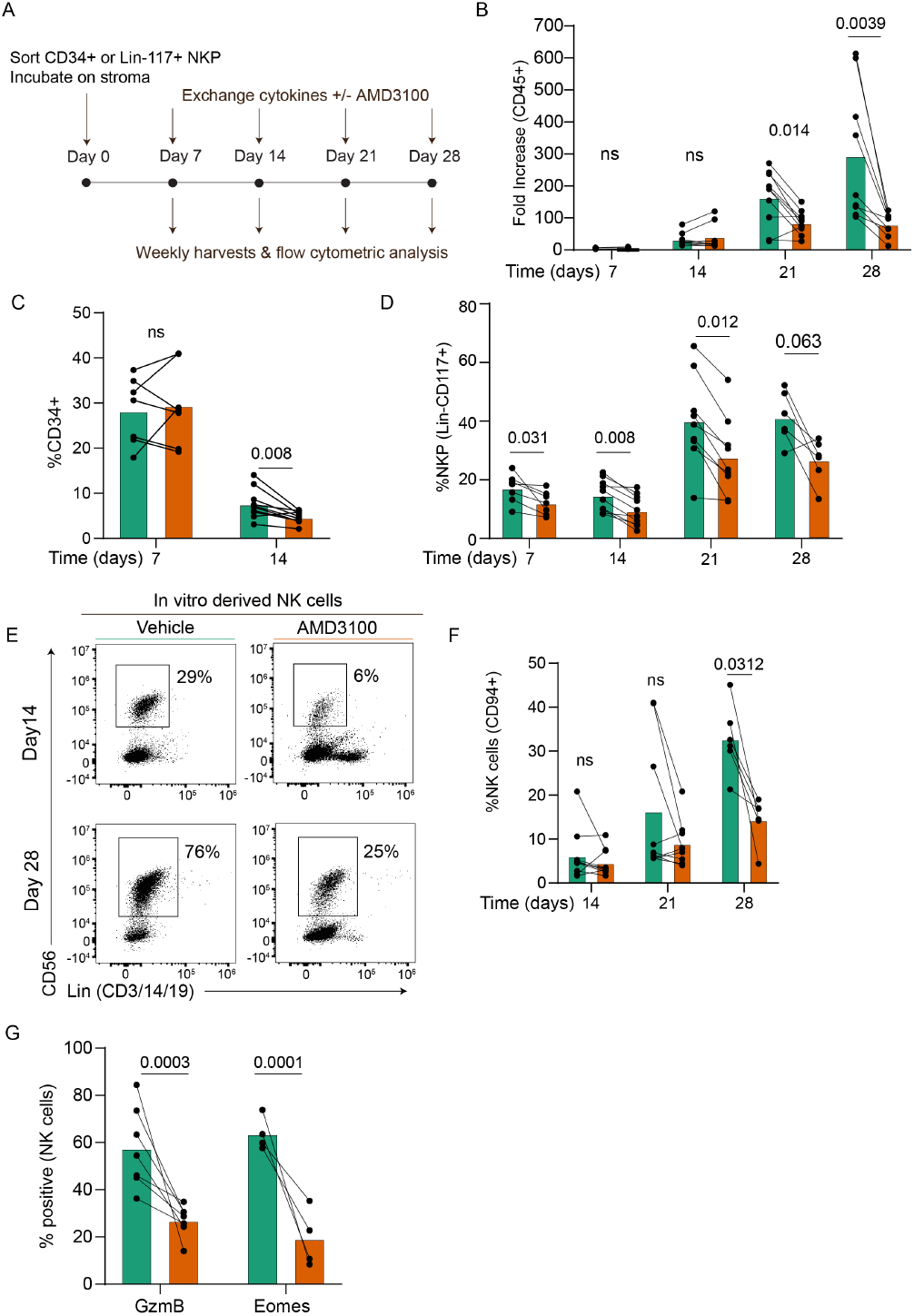
CXCR4-CXCL12 interactions are required for effective NK cell differentiation from CD34^+^ cells. A) Schematic of NK cell in-vitro differentiation protocol. Human CD34+ cells were isolated from cord blood using density centrifugation and fluorescence activated cell sorting (FACS). Sorted CD34^+^ cells were plated on EL08.1D2 stromal cells with exogenous cytokines as indicated. Cultures were treated with 10 µM AMD3100 or Vehicle at weekly media exchanges. Cells were harvested weekly for flow cytometric analysis. B) Fold increase in the absolute number of live CD45^+^ cells from AMD3100 (orange) and Vehicle (green) treated cultures. N = 4-10 donors. C) Frequency of CD34^+^ cells gated on CD45^+^ cells at days 7 and 14. n = 7-10 donors. D) Frequency of Lin^−^CD117^+^ NK/ILC progenitors gated on CD45^+^ live cells by week. n=7-9 donors. For B-D, each dot represents an individual donor and lines connect paired samples. Statistical comparisons between Vehicle and AMD3100 conditions at each time point were performed using Wilcoxon matched-pairs signed-rank tests. ns=not significant. E) Representative flow plots from days 14 and 28 showing the frequencies of CD56^+^ Lin^−^ NK cells. Previously gated on CD45^+^ live cells. F) Frequencies of CD94^+^ conventional NK cells at days 14, 21, and 28. n=7-9 donors. G) Frequencies of Granzyme B (GzmB) and EOMES positive NK cells at day 28. n=4-7 donors.

We first examined the fold increase of live CD45^+^ cells in each culture condition over time (Fig. 5B). Differences between vehicle and AMD3100-treated cultures were not significantly different through day 14. However, by day 21, a clear divergence emerged whereby vehicle-treated cultures underwent a robust expansion and AMD3100-treated cultures failed to expand. This indicates that CXCR4 signaling becomes particularly important during later stages of in-vitro NK cell differentiation when cells normally proliferate in response to stromal cues.

To test whether reduced output reflected impaired hematopoietic progenitor maintenance or differentiation, we measured CD34^+^ frequencies at days 7 and 14 and found no significant difference between conditions at day 7 (Fig. 5C), suggesting that early differentiation is not stalled in the presence of AMD3100. As expected, the frequency of CD34^+^ cells decreased from about 30% to under 10% between weeks 1 and 2 in both groups, consistent with CD34 down-regulation during progression through differentiation, and at day 14 we found significantly fewer CD34^+^ progenitors in the AMD3100-treated condition (10). Next, we looked at CD117^+^ Lin^−^ NK/ILC precursors (32), a substantial population of cells in-vitro differentiation cultures (8, 9). AMD3100 cultures contained significantly fewer NKPs than vehicle controls at multiple time points (Fig. 5D). This reduction in intermediate precursors suggests that CXCR4 signaling supports the progression and/or maintenance of developing NK cells beyond early progenitor stages.

By day 28, AMD3100-treated cultures showed a clear reduction in CD94^+^CD56^+^ mature NK cells (Fig. 5E, F). This observation was consistent with the failure of AMD3100 cultures to undergo the late expansion observed in vehicle-treated cultures. Representative flow cytometry plots illustrate the robust accumulation of CD56^+^ NK cells in vehicle cultures compared to the reduced population in AMD3100-treated cultures (Fig. 5E). NK cells generated in AMD3100 cultures also showed significantly reduced frequency of granzyme B^+^ and EOMES^+^ cells at day 28 (Fig. 5G). These markers reflect acquisition of effector functions and transcriptional maturation, indicating that CXCR4 blockade disrupts terminal NK cell differentiation.

In summary, AMD3100 treatment during in-vitro differentiation limits the generation and maturation of NK cells from CD34^+^ progenitors—reducing NK/ILC precursors and committed NK output and decreasing cytotoxic and transcriptional maturation markers. These findings suggest that CXCR4-CXCL12 signaling provides critical, stage-specific developmental cues—likely through niche positioning and stromal contact interactions—that support efficient human NK cell development.

### Plerixafor treatment of WHIM syndrome patients alters NK cell motility, mirroring in-vitro AMD3100 effects

Because our in-vitro data suggests that CXCR4 blockade alters NK cell phenotypes, we asked whether these features are mirrored ex-vivo in WHIM patient samples with gain-of-function CXCR4 mutations undergoing treatment (46). To characterize NK cell phenotype in WHIM syndrome, we analyzed peripheral blood from healthy donors (HDs) and WHIM patients by flow cytometry. All WHIM samples were collected following 8 months of treatment in a 12-month course of filgrastim (G-CSF) or the CXCR4 antagonist plerixafor (AMD3100) (Fig. 6A). Filgrastim mobilizes CXCR4^+^ cells by altering the CXCL12 in the stromal niche, while plerixafor directly blocks CXCR4-CXCL12 interactions (46). Since paired samples were available, we compared phenotypes during filgrastim versus plerixafor treatment from the same individual. Pre-treatment samples were not available.

**Fig. 6.**
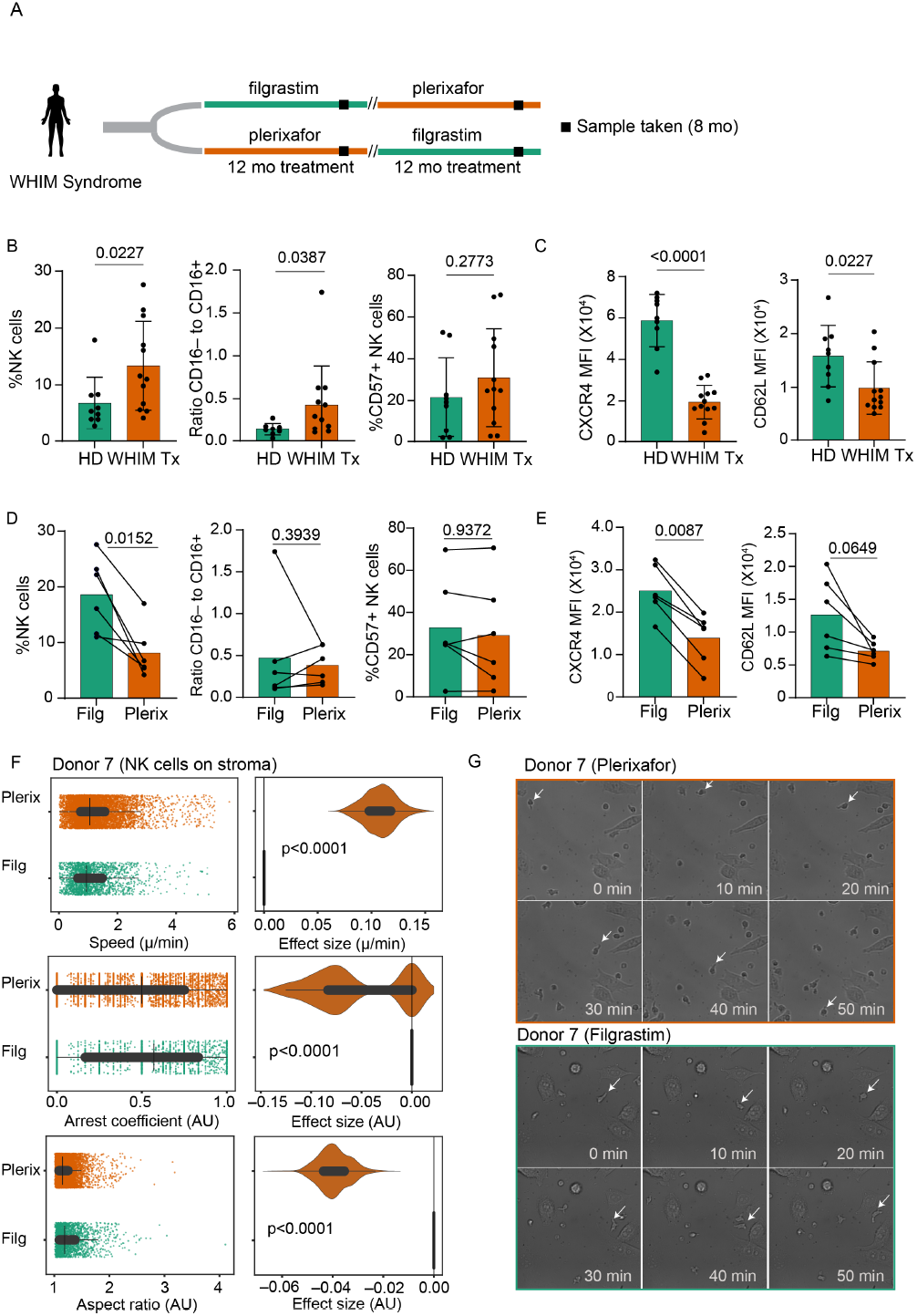
Plerixafor treatment of WHIM syndrome patients alters NK cell phenotype and motility. A) Schematic of WHIM patient sample collection and treatment timeline. All WHIM samples were obtained after 8 months of treatment following either filgrastim or plerixafor administration as previously described47. Paired samples from the same donor allowed for direct comparison of treatment effects. B) Flow cytometry quantification of NK cell frequency, CD16^+^ and CD57^+^ subsets in healthy donors (HDs) and WHIM patients treated with either filgrastim or plerixafor (WHIM Tx). NK cells were identified as CD45^+^CD3^−^CD14^−^CD19^−^CD56^+^ lymphocytes. n=9 HD samples and 12 WHIM patient samples. Data shown as mean±SD; p-values calculated by Mann-Whitney test. C) Surface CXCR4 and CD62L expression on NK cells from HD and treated WHIM patients (WHIM Tx); mean±SD; p-values calculated by Mann-Whitney test. D) NK cell frequency, CD16^+^ and CD57^+^ subsets in paired samples from the same individual undergoing filgrastim (Filg) or plerixafor (Plerix) treatments. n=6 donors undergoing each treatment; mean±SD; statistical significance determined using Wilcoxon matched-pairs signed-rank tests.. E) Surface CXCR4 and CD62L expression on NK cells from paired samples from the same individual undergoing filgrastim or plerixafor treatments. n=6 donors undergoing each treatment; mean±SD; p-values calculated by Mann-Whitney test. F) NK cells were isolated from peripheral blood of WHIM syndrome individuals undergoing plerixafor or filgrastim treatment and incubated on EL08.1D2 stromal cells then imaged by confocal microscopy for 4 hours at 2 min intervals. Quantified motility and shape parameters comparing NK cells of treated WHIM patients. Data are paired between post-filgrastim and post-plerixafor samples from twelve WHIM donors. Point plots display pooled single-cell metrics with effect size distributions. G) Representative time-lapse imaging of post-plerixafor and post-filgrastim treated NK cells (white arrows) on stromal cells.

WHIM patients displayed a significantly increased percentage of NK cells among live CD45^+^ cells from peripheral blood compared to HDs (Fig. 6B). NK cells from WHIM patients also showed a skewing toward the less mature CD16^−^ population, whereas the proportion of terminally differentiated CD57^+^ NK cells did not differ significantly between groups (Fig. 6B). This immature bias may reflect altered differentiation or trafficking dynamics associated with CXCR4 dysregulation and/or mobilizing therapies.

Given their central roles in NK cell trafficking and tissue localization, we next examined CXCR4 and CD62L expression. Surface CXCR4 expression was modestly lower on NK cells from treated WHIM patients compared to HDs (Fig. 6C). Given that CXCR4 gain-of-function mutations in WHIM are classically associated with impaired receptor internalization and heightened CXCL12 responsiveness, this is surprising. However, this reduction in detection likely reflects the residual binding of plerixafor to CXCR4, which is known to compete for antibody binding with detection antibody. Additionally, CD62L expression was significantly reduced in WHIM patient NK cells (Fig. 6C), and this reduction was seen in both CD16^−^ and CD16^+^ subsets (not shown), indicating that the difference is not driven by changes in subset composition.

To more directly analyze treatment effects, we compared paired samples during filgrastim or plerixafor treatment from individual WHIM donors. The frequency of CD56^+^ NK cells among CD45^+^ leukocytes was lower after plerixafor treatment compared to filgrastim (Fig. 6D). Moreover, CXCR4 surface levels were lower in plerixafor-treated NK cells relative to filgrastim-treated samples (Fig. 6E), consistent with plerixafor occupancy blocking the 12G5 antibody epitope. CD62L expression also trended lower under plerixafor treatment, though this difference did not reach significance.

To assess how treatment influences NK cell migratory behavior, CD56^+^ NK cells were sorted from patient samples and plated on EL08.1D2 stromal monolayers. NK cells from filgrastim versus plerixafor treated WHIM patients displayed striking differences in morphology and motility (Fig. 6F, G). NK cells from filgrastim-treated patients showed more elongated and polarized morphologies, whereas NK cells from plerixafor-treated patients were markedly rounded. Additionally, NK cells from plerixafor-treated patients exhibited lower aspect ratio, reduced elongation, and increased rounding (Fig. 6F), indicating impaired polarization. They also showed significantly lower arrest coefficient, suggesting reduced stable substrate engagement. Instantaneous speeds were modestly higher in NK cells from plerixafor-treated patients, consistent with a non-adherent, exploratory mode of movement. Examples from time-lapse imaging confirmed these differences, as filgrastim-treated NK cells formed persistent leading edges and stable contacts with stroma, whereas plerixafor-treated NK cells remained spherical, transiently contacting the surface but failing to establish durable adhesions (Fig. 6G). This phenotype closely mirrors the reduced adhesiveness and polarization we observed with AMD3100 (plerixafor) treatment in-vitro, reinforcing that CXCR4 blockade directly disrupts adhesion and cytoskeletal organization.

Taken together, these findings indicate that pharmacologic CXCR4 antagonism disrupts NK cell polarization and stable substrate engagement, potentially impairing their ability to navigate tissues or form long-lived immune synapses. While plerixafor is clinically effective for leukocyte mobilization and WHIM management, our results suggest that transient impairment of NK cell adhesion and polarity may represent a short-term tradeoff of CXCR4 blockade, with potential implications for NK surveillance and effector functions during treatment.

## Discussion

Our data support a model in which CXCR4-CXCL12 interactions orchestrate NK cell positioning and adhesion within developmental niches, thus enabling efficient NK cell maturation. CXCL12 is highly expressed on tonsil stromal cells, particularly on fibroblastic reticular cells (47, 48), and is abundant in para- and interfollicular regions that also support developing NK cells. In the thymus, CXCL12 is crucial for positioning T cells for effective T cell development and proliferation, and in the bone marrow, CXCL12-associated reticular (CAR) cell niches are sites of NK cell development (16, 49, 50). As fibroblastic reticular cells in lymph node can also produce IL-15 (51), which is necessary for NK cell maturation, we propose that CXCL12 could play a role in retaining HSPCs and NKPs in sites that are enriched with cytokines that promote NK cell differentiation. This model is analogous to the role of CARs in murine bone marrow (16) and suggests that CXCR4-CXCL12 signaling integrates tissue localization with access to differentiation cues.

Beyond positioning NK cells within CXCL12-rich niches, our data indicate that CXCR4 plays a key role in balancing NK cell motility and adhesion. Pharmacologic blockade of CXCR4 results in cells that are more motile, less arrested, and rounder on developmentally supportive stromal cells. Consistent with reduced adhesion, CXCR4 inhibition impairs β2 integrin activation, suggesting that CXCR4 stabilizes cell contacts. CXCR4-dependent adhesive structures have been described in other hematopoietic lineages; HSPCs form highly polarized, CXCR4-CXCL12-dependent “magnupodia” that anchor them to bone marrow stromal cells (15, 52). Developing T cells at the β-selection checkpoint likewise assemble a CXCR4-dependent immunological synapse with thymic stroma that is required for progression through this developmental checkpoint (53, 54). This CXCR4-dependent shift in cells from an adhesive, elongated migration phenotype to a rounded, high motility state demonstrates how NK cells can integrate environmental cues to transition between migratory and adherent behaviors. Although this balance is classically described in the context of a cytotoxic synapse (55), a similar principle likely applies in non-cytolytic developmental settings. CXCR4-CXCL12 signaling provides positional information that coordinates adhesion, integrin activation, and motility to regulate how NK cells engage with stromal cells and extracellular matrix.

The functional consequences of this altered adhesion are evident in our differentiation system, where AMD3100-treated cultures show reduced generation of mature NK cells and incomplete cytotoxic maturation. These observations are consistent with prior work showing that NK differentiation on EL08.1D2 stroma is contact-dependent (11). In addition, our previous description of the developmental synapse demonstrated that developing NK cells polarize and form stable CD56 and actin-rich contacts with stromal cells are required for efficient human NK cell differentiation (12). These findings suggest that reduced stable interactions with stromal cells under CXCR4 blockade prevent progenitors from establishing polarity or stable signaling contacts with stroma that are required for full NK lineage commitment.

The relevance of these findings is underscored by our analysis of samples from patients with WHIM syndrome, a disorder driven by hyperactive CXCR4 signaling (25). Although CXCR4 dysfunction in WHIM syndrome has been well studied in neutrophils and B cells, its impact on NK cells has been less clear, with prior reports describing either reduced or unchanged NK cell frequencies in peripheral blood (26, 27). In our cohort, NK cells collected during plerixafor treatment exhibit a rounded morphology and reduced arrest behavior, closely mirroring the phenotype of AMD3100-treated NK cells in-vitro. Detection of surface CXCR4 and CD62L were also reduced during plerixafor treatment, consistent with CXCR4 receptor occupancy and altered trafficking. These findings suggest that while pharmacologic CXCR4 antagonism is effective for hematopoietic mobilization, it may transiently impair the ability of NK cells to form polarized, adhesive contacts with substrates. It is important to note that filgrastim and plerixafor mobilize leukocytes through distinct mechanisms, namely a G-CSF-mediated remodeling of the stromal niches versus direct CXCR4 antagonism (26, 56), which likely contributes to the treatment-dependent differences in NK behavior we observed. Additionally, as expected, differences between in-vitro and ex-vivo phenotypes likely reflect distinct drug exposure dynamics.

Several limitations should be considered when interpreting these findings. First, our WHIM cohort size was limited, and untreated baseline samples were not available, preventing direct assessment of the intrinsic NK phenotype in WHIM syndrome. Second, this study overall did not examine potential crosstalk between CXCR4 and other receptors, including S1P5, which is known to influence NK cell egress58 and may intersect with CXCR4-dependent positioning. Third, although EL08.1D2 stroma recreate key features of CXCL12-rich niches, in-vivo developmental contacts likely involve additional stromal subsets and extracellular matrix components not modeled here. Nonetheless, the convergent in-vivo and in-vitro observations support a model in which CXCR4 signaling regulates NK cell adhesion and motility in human physiology. Supporting this broader role, disruptions of the CXCR4-CXCL12 axis in tumor models enhances NK cell infiltration (57), also mirroring the increased migration we observe under CXCR4 blockade.

Together, these data identify CXCR4-CXCL12 as a key regulator of the balance between migration, adhesion, and maturation in human NK cells. Our findings place CXCR4-CXCL12 at the center of a regulatory circuit that coordinates NK cell adhesion, motility, and developmental progression. By tuning integrin activation and stabilizing interactions with stromal niches, CXCR4 helps ensure that developing NK cells receive the signals necessary for efficient maturation.

## Methods

### Sex as a biological variable

Sex was not considered as a biological variable in this study. Human peripheral blood and umbilical cord blood samples were obtained from de-identified donors, and donor sex information was not available. As a result, samples were included without sex-based stratification, and analyses were focused on CXCR4–CXCL12–dependent cellular behaviors across donors rather than sex-specific differences.

### Public dataset analysis

CXCR4 expression data for circulating lymphoid progenitors and NK cell subsets were obtained from the Human Immune Health Atlas (29). For each annotated cluster, we calculated mean CXCR4 transcript levels and the proportion of CXCR4-expressing cells. Bubble plots were generated in Python using Scanpy for data processing and Matplotlib to visualize average expression (color scale) and frequency of expressing cells (bubble size).

### Study approval and sample processing

All human samples were collected in accordance with the Declaration of Helsinki and approved by the Columbia University or NIH Institutional Review Boards (IRB). Informed consent was obtained from all donors or their legal guardians prior to sample collection and all samples were de-identified prior to analysis.

Peripheral blood was obtained from healthy donors via venipuncture at Columbia University Medical Center. Tonsil samples were acquired from routine tonsillectomies on pediatric patients at Columbia University Medical Center. Tonsil tissue was mechanically dissociated by mincing with sterile scalpels and passing the suspension through a 40-µm cell strainer (Falcon, 352340). Cell suspensions were washed with PBS by centrifugation at 300xg for 10 minutes at room temperature. For flow cytometric analysis, cells were resuspended in PBS and stained with relevant antibodies or resuspended in FBS 10% DMSO freezing media and cryopreserved.

De-identified PBMCs were isolated from the blood of participants in a clinical trial of the treatment of WHIM patients with plerixafor versus filgrastim (46) (Clinicaltrials.gov NCT02231879). Cryopreserved cells from participants were deidentified and used in this study. The protocol was approved by the National Institute of Allergy and Infectious Diseases Institutional Review Board and all participants signed informed consent prior to participation. This was a single center, investigator-initiated, blinded cross-over trial where each participant received each treatment in succession with a randomization of which drug started first. Both drugs were administered twice daily subcutaneously using prefilled syringes and matched samples used here were drawn at the 6-month time point of the one-year treatment period (8 months of drug exposure counting a 2-month initial dose adjustment period). Untreated baseline samples were not available from these donors.

De-identified umbilical cord blood samples were acquired from the placenta of women who were undergoing routine healthy term deliveries (between 37 weeks 0 days and 41 weeks 6 days gestation). Blood was collected in either BD Vacutainer with Sodium Heparin or BD Vacutainer with 7.2mg K2 EDTA. Collection followed institutional SOPs and was performed in accordance with the AIDS Clinical Trials Group and IMPAACT Network Umbilical Cord Blood Collection Standard Operating Procedure (58).

Mononuclear cells were isolated from whole blood and cord blood by Ficoll-Paque (Cytiva; catalog no. 17144003) density gradient centrifugation. NK cells and progenitor cells were enriched using StemCell Rosette Sep negative selection enrichment kits (NK cell enrichment, STEMCELL technologies; catalog no. 15065; Hematopoietic stem cell enrichment, STEMCELL technologies; catalog no. 15066) following manufacturer instructions.

For in-vitro differentiation experiments, CD34^+^ precursors were isolated from cord blood samples. Mononuclear cells isolated from donors were incubated with anti-CD34 antibody (Biolegend; Clone: 581) and CD34^+^ cells were isolated by FACS on Sony MA900 sorter.

### Cell culture

NK92 cells (ATCC) were maintained in complete NK92 growth medium consisting of α-MEM without ribonucleosides or deoxyribonucleosides (Gibco 12-561-056) supplemented with 12.5% heat-inactivated horse serum, 12.5% heat-inactivated FBS, 2mM L-glutamine, 1.5 g/L sodium bicarbonate, 0.2 mM myo-inositol, 0.1 mM β-mercaptoethanol, 0.02 mM folic acid, and 100 U/mL recombinant human IL-2. Medium was filtered prior to use. NK92 cells were maintained at 37°C with 5% CO2 and passaged every 2-3 days. EL08.1D2 stromal cells (provided by E.A. Dzierzak, Erasmus MC, Netherlands, via J.S. Miller, University of Minesota) were cultured on gelatin-coated plates at 32°C in medium composed of 40.5% α-MEM (Life Technologies), 50% Myelocult (STEMCELL Technologies), 7.5% heat-inactivated FBS supplemented with 2mM GlutaMAX, 100U/mL penicillin/streptomycin, 50mM β-mercaptoethanol, and 10^−6^M hydrocortisone. For co-culture experiments, EL08.1D2 cells were plated into gelatinized 96-well plates and mitotically inactivated by irradiation at 300 rads prior to the addition of progenitor cells.

### Flow cytometry

Freshly isolated or cryopreserved cells were thawed into warm R10 media and then washed with PBS. Cryopreserved cells were allowed to recover in R10 media at 37°C for 3 hours before staining or sorting. Cells were stained for extracellular markers for 30 minutes at 4°C and then washed with PBS. When intracellular markers were stained, cells were fixed and permeabilized using eBioscience FoxP3 buffer (catalog no. 00-5523-00; ThermoFisher) and then stained for intracellular markers for 30 minutes at 4°C. Data was acquired on a Novocyte Penteon Cell Analyzer and then exported to FlowJo (BD Biosciences) for analysis. Antibodies used for flow cytometry are provided in Supplemental Table 1.

### Transwell migration assay

Transwell migration assays were performed using Corning Transwell permeable supports with a 6.5mm insert, 8.0 µm pores in a 24 well plate (catalog no. 3422) or CELLTREAT permeable cell culture inserts in 12 well plate, 8.0 µm pores (catalog no. 230619). For each condition, 600 µl Opti-MEM media (Thermo Fisher) supplemented with CXCL12 (200 ng/mL) and/or AMD3100 (10 µM) was added to the lower chamber. Recombinant IL-2 (100 U/mL) was included for NK92 cells. Cells were pelleted, resuspended in Opti-MEM containing IL-2±AMD3100, and 5 x 10^5^ cells in 100 µl were added to the upper chamber. Assays were performed in duplicate. Plates were incubated for 5h at 37°C, and migrated cells were collected by rinsing the underside of each insert with 400 µl PBS. Cells in the lower well were counted.

### CRISPR-CAS9 gene editing

Two million NK92 cells were nucleofected with 5 µM of sgRNA guide specific for CXCR4 (Horizon; SgRNA CXCR4 (ATGCTGATCCCAATGTAGTA); Cat SG-005139-01-0002) and 5 µg of Cas9-GFP mRNA (Horizon) in 100 µL of nucleofection Kit-R supplemented (Lonza) solution. The R-024 program of an Amaxa Nucleofector II (Lonza) was used. After 24h GFP+ cells were sorted and cultured prior to validation.

### Cytotoxicity assay

Four-hour Cr51 release assays were performed using K562 target cells as described previously (59). Target cells were used at a concentration of 10^4^ target cells per well of a 96-well U bottom plate. Effector cells were used in serial dilution starting at an effector to target cell ratio of 50:1 for PBMC or 10:1 for NK92 cells. After incubation, supernatants were transferred into Lumaplates (Revvity) and measured using a TopCountXL (PerkinElmer). Specific lysis (%) was calculated as: (experimental cpm – spontaneously released cpm)/(total cpm – spontaneously released cpm) ×100.

### Cyclic immunofluorescence analysis of FFPE pediatric tonsil sections

Formalin fixed parafilm embedded (FFPE) tonsil sections were stored at -20°C and brought to room temperature before processing. After three 10-minute Xylene (catalog no. X5–1; Fisher Scientific) washes, tissue was rehydrated in by sequential incubations in 100% and 95% ethanol (catalog no. BP2818100; Fisher Scientific) and deionized (DI) water for 10 minutes, and then in 70% and 35% ethanol and DI water for 5 minutes. Heat-activated antigen retrieval was performed by incubating slides in pre-warmed Agilent Dako pH6 antigen retrieval solution (catalog no. S2367; Agilent) for 20-minutes in a vegetable steamer (Black Decker), cooled to room temperature, and washed with PBS.

Tissue sections were incubated with 1 mg/mL borohydride in deionized water for 10 minutes to decrease background fluorescence, washed three times with PBS, permeabilized with 0.2% Triton X-100 in PBS for 30 minutes, and washed with PBS three times. Next, a hydrophobic border was drawn with a PAP pen and tissue samples were incubated with ThermoFisher Super Blocking Buffer (catalog no. 37580; ThermoFisher) for 1 hour.

Antibodies for immunostaining (Supplemental Table ) were diluted in an antibody diluent buffer (PBS/10% Bovine Serum Albumin/0.01% Triton X-100). Immunostaining of tissue sections with unconjugated antibodies for high-resolution immunofluorescence microscopy was done overnight at 4°C in a humidity chamber. Slides were washed with antibody diluent buffer four times after immunostaining. Tissue sections were incubated with secondary antibodies at a dilution of 1:200 for 1 hour at room temperature followed by two washes with antibody diluent buffer and five washes with PBS. PBS was removed, then tissue was treated with Vector TrueVIEW Autofluorescence Quenching Kit (catalog no. SP-8400; Vector Labs) prior to mounting. After washing excess TrueView reagents three times with PBS, tissue sections were mounted using VectaShield Antifade Mounting Medium with DAPI (catalog no. H-1200; Vector Laboratories, Inc) and a 1.5 coverslip (catalog no. 72204-03; Electron Microscopy Services). Regions of interest were imaged with a 20% overlap using a Zeiss CellDiscoverer 7 using a 20X/0.7 or 20X/0.95 NA objective and an optical magnification of 0.5X and coordinates were saved for subsequent imaging. This resulted in images with a micron per pixel unit of 0.454 microns/pixel. After imaging, coverslips were gently removed by washing with PBS in coplin jars.

To strip antibodies, tissue slides were incubated in freshly prepared stripping buffer (20 mL 10% SDS, 0.8 mL beta-mercaptoethanol, 12.5 mL 0.5 M Tris-HCl (pH 6.8), 67.5 mL DI H2O) for 40 mins in a vegetable steamer then washed with water for 15 mins. Slides were then blocked with SuperBlock for 10 minutes and re-stained with primary and secondary antibodies prior to re-imaging as described above. Tile stitching, channel registration, and barrel-distortion correction were performed using Ashlar (60). All downstream visualization was performed in Fiji (61).

### Proliferation assays

NK92 WT and NK92 CXCR4-KO cells were cultured in complete NK92 medium at 37°C and 5% CO2. Cells were seeded at 2 x 10^5^ cells/mL in 24-well plates and treated with AMD3100 (10 µM) or vehicle (water) control. Live cell concentrations were measured at 0, 24, 48, and 72 hours using AOPI staining and an automated cell counter (Cellometer).

### Confocal imaging and data analysis

For fixed-cell imaging of stromal cells, EL08.1D2 stromal cells were cultured on gelatin-coated chamber slides (Ibidi) until 70–80% confluent. Cells were fixed with 4% paraformaldehyde for 10 minutes at room temperature, washed with PBS, and permeabilized with 0.1% Triton X-100 for 10 minutes. After blocking with 5% BSA in PBS for 1 hour, cells were incubated with anti–CXCL12 conjugated antibody (clone 79018, RnD, FAB350T, 1:100) and Phalloidin (Thermofisher, A22287, 1:200) for 1 hour at room temperature. Slides were washed three times in PBS, followed by DAPI nuclear staining.

Primary NK cells were washed and incubated in HBSS with Ca^2^Mg^2^ for 30 min at 37°C. AMD3100 (10 µM) was added for 30 min where indicated. Cells were stimulated with CXCL12 (200 ng/mL) for 5 min and immediately fixed in 4% paraformaldehyde for 10 min. After washing and permeabilizing with 0.1% Triton X-100, cells were stained with the anti-LFA-1 Alexa Fluor 488 antibody (clone m24, Biolegend, 363404), phalloidin Alexa Fluor 568 (Thermo A12380), and DAPI (Millipore, D9542).

Confocal images were acquired on a Yokogawa spinning disc confocal on a Zeiss Axioimager Z1 stand with a 63x 1.4 NA objective described for live cell imaging using matched laser and exposure settings. Images were processed and analyzed using Slidebook (3i) and Fiji (ImageJ) (61). Z-stacks (0.3–0.5 µm steps) were collected and maximum-intensity projections were generated in Fiji. Cell segmentation was performed using actin masks and m24 MFI and m24 membrane area measurements were extracted for each cell. At least 10 cells per condition per donor were analyzed.

### Live-cell migration imaging and analysis

Primary NK cells, CD34^+^ progenitors, or NK cell lines were pretreated with vehicle control or AMD3100 (10 µM) for 2 hours at 37°C. For stromal co-cultures, EL08.1D2 stromal cells were plated as confluent monolayers 24 hours before imaging. For integrin ligand surfaces, wells were coated overnight at 4°C with recombinant human ICAM-1 Fc or VCAM-1 Fc (5 µg/mL), followed by two PBS washes. Cells were added gently to the monolayer or ligand surface in NK media. Live-cell imaging was performed on a spinning-disk confocal microscope equipped with a 37°C 5% CO environmental chamber. Brightfield images were acquired every 2 minutes for 2–6 hours using a 20× objective on a Yokogawa spinning disc confocal on a Zeiss Axioimager Z1 stand with a 63x 1.4 NA objective. Time-lapse movies were exported as TIFF stacks for analysis.

Segmentation of individual cells was performed using Cellpose (v2) (39). Segmentation masks were visually inspected for accuracy. Cell tracking was performed using btrack (40). Tracks shorter than 2 frames were removed. Tracking results were visually reviewed in napari to confirm accuracy. Segmentation and tracking outputs were filtered, as described for each experiment.

Quantitative migration metrics were computed using cellPLATO as described by Shannon et al. (41), including cumulative track length, instantaneous speed, and arrest coefficient (fraction of frames in which displacement was <1 µm/min). Morphological features (cell area, aspect ratio, and solidity) were calculated from segmentation masks on a perframe basis and averaged for each cell.

Migration and shape metrics were analyzed at the single-cell level. Differences between treatment conditions were evaluated using bootstrap resampling and permutation-based P values. Effect size and confidence intervals were visualized using plots of differences. Statistical analyses were performed using custom Python scripts.

### In-vitro NK cell differentiation on EL08.1D2 stromal cells

In-vitro NK cell differentiation was performed as previously reported (12). Human CD34^+^ hematopoietic progenitors were isolated from cord blood by FACS sorting and 400 cells per well were plated onto an irradiated EL08.1D2 stromal cell monolayer. Differentiation cultures were maintained in Ham F12 media plus DMEM (1:2) with 20% human AB serum, ethanolamine (50 µM), ascorbic acid (20 µg/ml), sodium selenite (5 ug/l), β-mercaptoethanol (24 µM), and penicillin/streptomycin (100 U/ml) with IL-15 (5 ng/ml), IL-3 (5 ng/ml), IL-7 (200 ng/ml), SCF (20 ng/ml), and Flt3L (10ng/ml). Half of the media volume was exchanged weekly. For AMD3100 conditions, 10 µM AMD3100 was added to cultures with weekly media exchanges. Developing NK cells were collected at indicated timepoints for flow cytometry analysis.

### Statistics

Statistical analyses were performed with Prism 10.0.0 (GraphPad). Unpaired samples were measured with parametric (two-tailed t-test) or non-parametric (Mann-Whitney) tests depending on whether data fell within a normal distribution. Paired data were compared using Wilcoxon paired ranked-signs test. Paired comparisons across NK developmental populations were analyzed using a Friedman test with Dunn’s correction for multiple comparisons. Each donor was treated as an independent biological replicate. Outliers were identified using the ROUT method (Q = 1%) in GraphPad Prism.

## Data availability

Supplemental Movies 1-12 are available here: https://doi.org/10.5281/zenodo.18986487.

## ACKNOWLEDGEMENTS

We thank members of the Mace laboratory for scientific discussion and technical assistance. Evelyn Almonte Hernandez, Tolani Aliyu, and Hijab Fatima provided critical assistance with acquiring tonsil tissue and peripheral blood samples from healthy donors. We acknowledge collaborators for providing samples and expertise, and the Columbia University Stem Cell Initiative for the flow cytometry core facility and specifically Michael Kissner for experimental support. We would also like to thank the Herbert Irving Comprehensive Cancer Center Histology Services and Tumor Banking core at Columbia University Irving Medical Center for assistance with tissue embedding and sectioning. We are grateful to the donors who contributed samples for this study. This work was partially funded by T32GM007088 to S.E. and R01AI137073 to EMM. The work on WHIM patient samples was partially funded by the Division of Intramural and Clinical Research of the National Institute of Allergy and Infectious Diseases.

