## Supplemental Figures 1-4 for "CXCR4 coordinates adhesion, migration, and development of human NK cells"

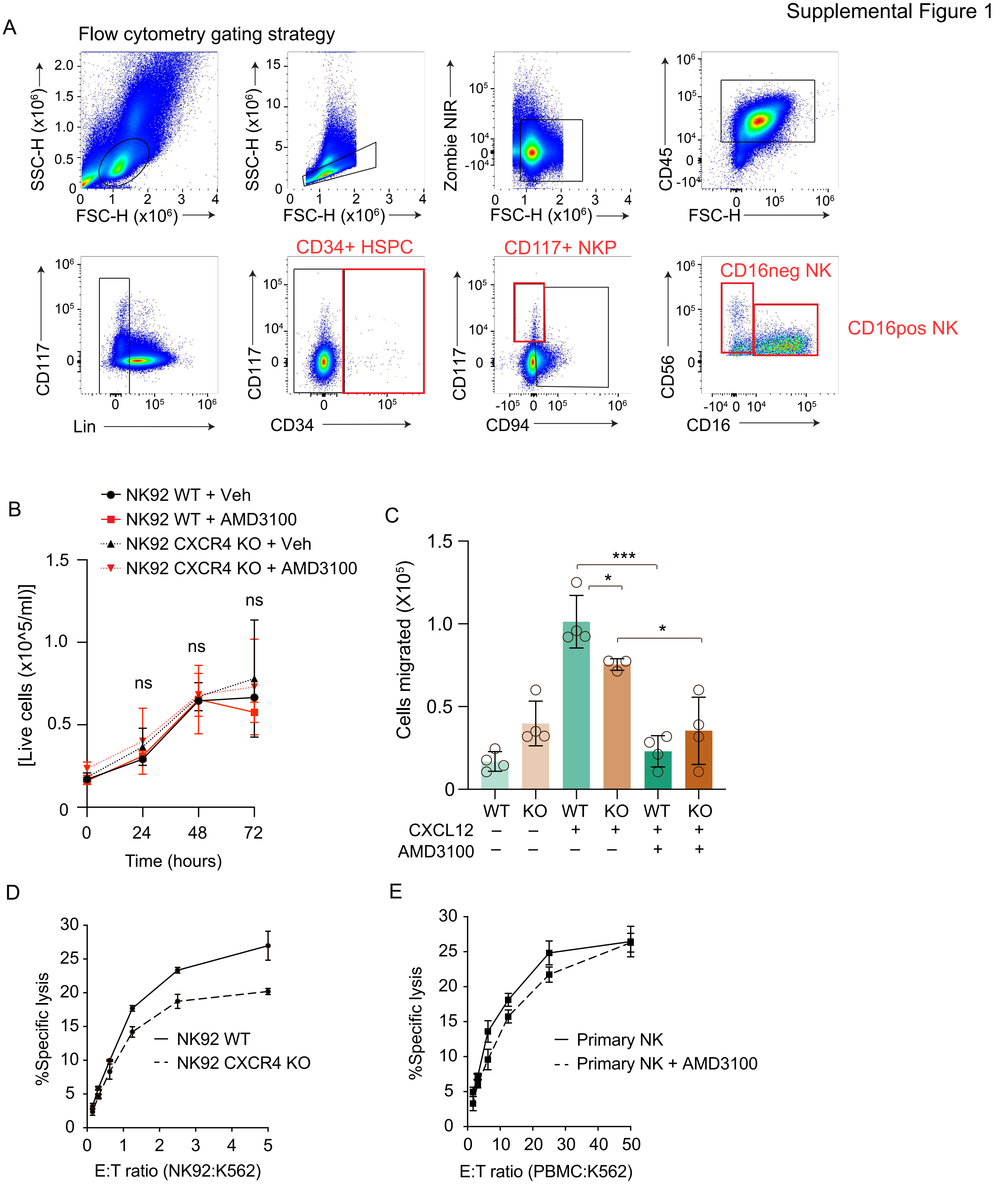


**Supplemental Figure 1. Gating strategy, proliferation assays, transwell assays, and killing assays** A) Gating strategy used for analysis of NK cell developmental intermediates in peripheral blood and tonsil. B) NK92 WT and CXCR4 KO cell proliferation measured over 72h in the presence of Vehicle (water) or 10μM AMD3100. Lines show mean±SD (n=3 independent experiments). ns=not significant by two-way ANOVA. C) CXCL12-directed chemotaxis of NK92 WT and CXCR4 KO cells ±AMD3100 in transwell migration assays; mean±SD (n=3 independent experiments). P values determined by Mann-Whitney test between samples as indicated. D) Chromium cytotoxicity assays using NK92 (left) or peripheral blood mononuclear cells (right) co-cultured with K562 target cells at indicated effector: target (E:T) ratios. Mean±SEM.

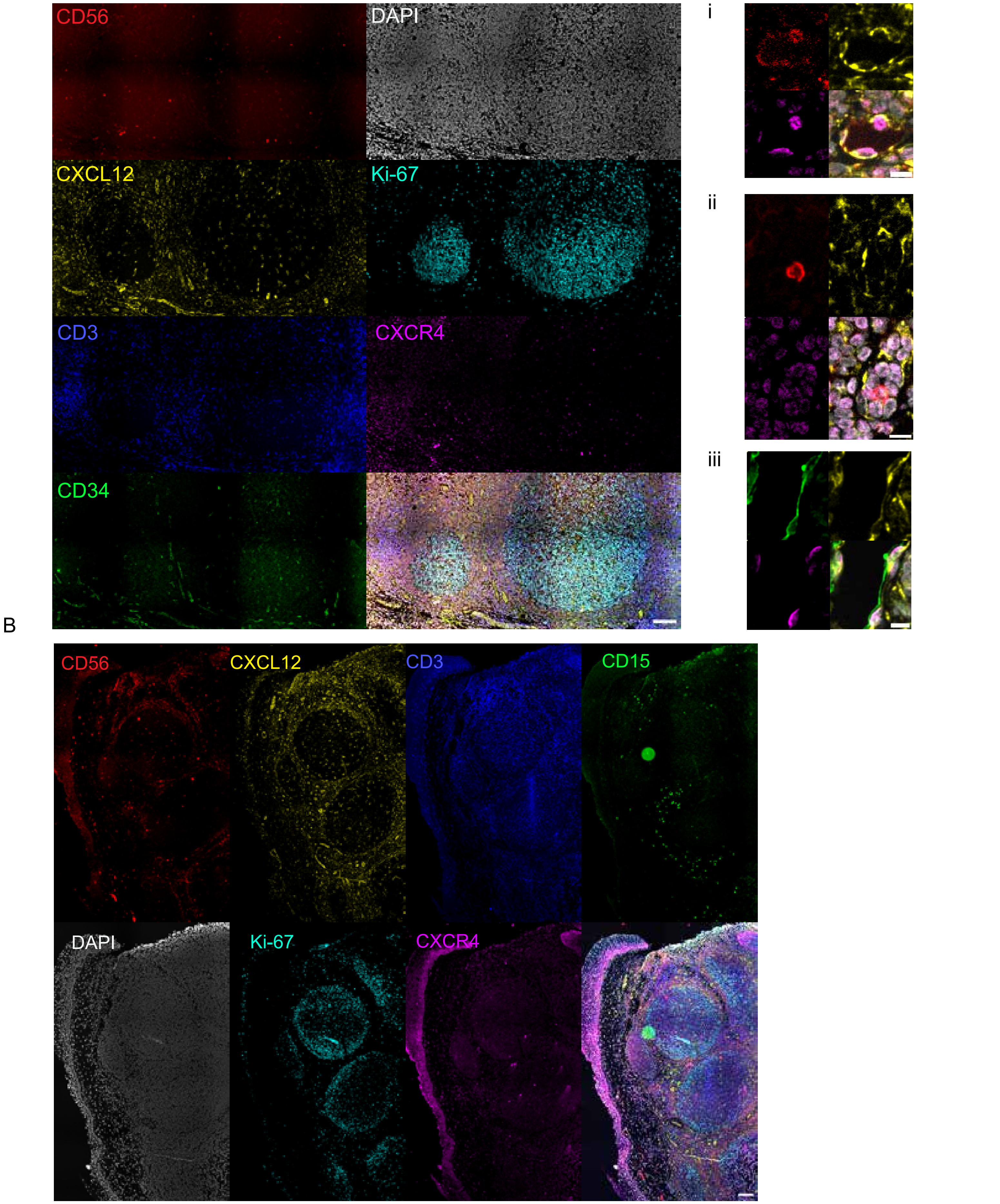


**Supplemental Figure 2.** Cyclic immunofluorescence (CyCIF) imaging of pediatric tonsil identifying CXCL12-rich stromal regions and associated immune populations. A) Representative CyCIF panel from pediatric tonsil tissue showing individual channels for CD56 (red), DAPI (gray), CXCL12 (yellow), Ki-67 (cyan), CD3 (blue), CXCR4 (magenta), and CD34 (green). Insets highlight CXCR4+ CD56+ and CXCR4+ CD34+ cells positioned near CXCL12-producing stromal structures. B) Second representative region with individual channels for CD56 (red), DAPI (gray), CXCL12 (yellow), Ki-67 (cyan), CD3 (blue), CXCR4 (magenta), and CD15 (green).

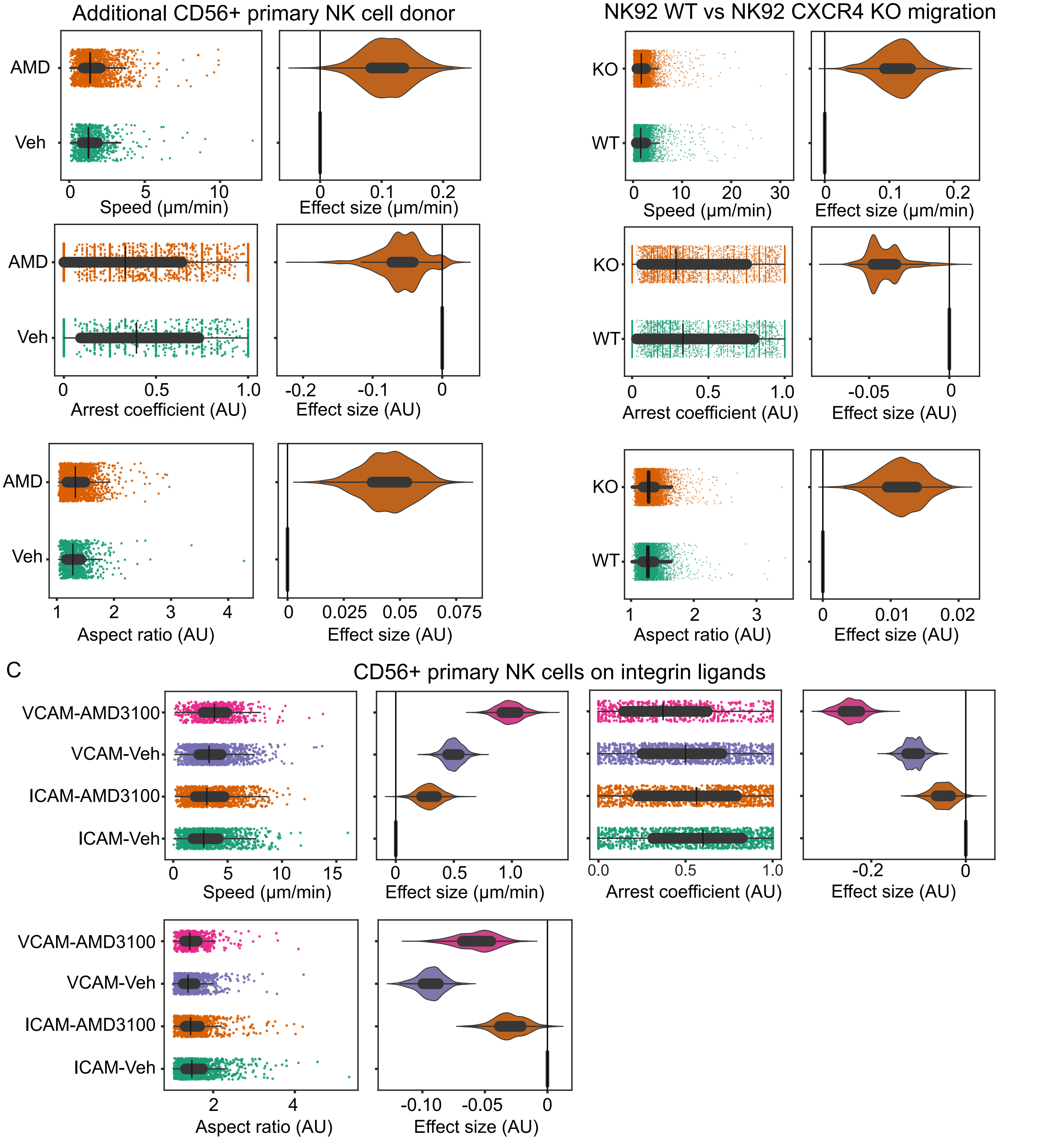


**Supplemental Figure 3. Plots of migration and morphological metrics from additional donors**. A) NK cells were isolated from peripheral blood of healthy donors and incubated on EL08.1D2 stromal cells in the presence of AMD3100 or Vehicle as indicated then imaged by confocal microscopy for 4 hours at 2 min intervals. Quantified motility and shape parameters comparing NK cells of treated WHIM patients. Point plots display pooled single-cell metrics with effect size distributions. B) NK cells were isolated from peripheral blood of healthy donors and incubated on recombinant VCAM1 or ICAM1 in the presence of AMD3100 or Vehicle as indicated then imaged by confocal microscopy for 4 hours at 2 min intervals. Quantified motility and shape parameters comparing NK cells of treated WHIM patients. Point plots display pooled single-cell metrics with effect size distributions.

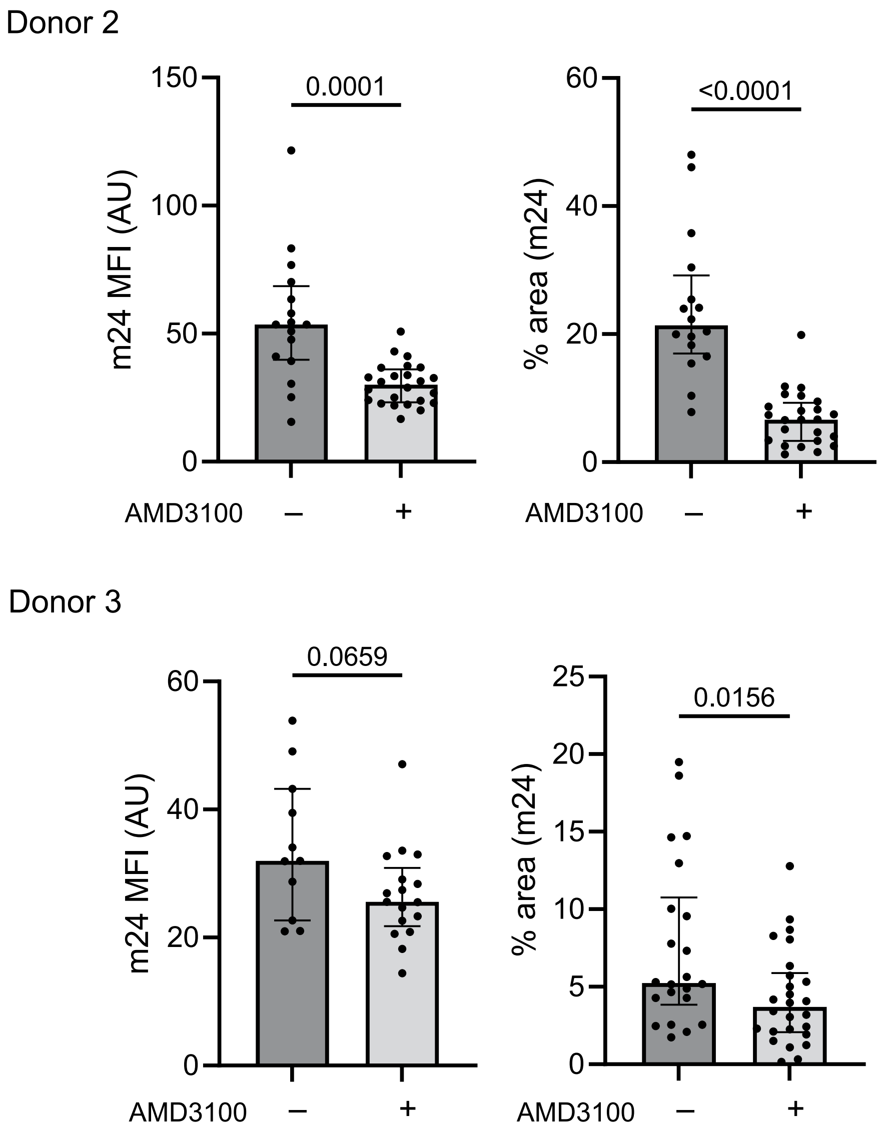


**Supplemental Figure 4.** Additional donors for m24 MFI and percent area. Quantification of m24 MFI and percent area from two additional healthy donors (see Fig. 4E for Donor 1). Each dot represents a single cell from one donor. Horizontal bars indicate mean±SD. Statistical comparison was performed using a two-tailed Mann-Whitney test.
