## Supplemental Tables for "CXCR4 coordinates adhesion, migration, and development of human NK cells"

***Supplemental Table 1: Antibody panel used for flow cytometric analysis of NK cell developmental subsets***

| ANTIGEN | CLONE | FLUOROCHROME | VENDOR | CATALOG # | DILUTION |
|---|---|---|---|---|---|
| **CD45** | HI30 | BUV805 | BD Biosciences | 612891 | 1:100 |
| **CD94** | HP-3D9 | BUV395 | BD Biosciences | 743954 | 1:150 |
| **CD3** | UCHT1 | BUV496 | BD Biosciences | 612940 | 1:200 |
| **CD14** | M5E2 | BUV496 | BD Biosciences | 750381 | 1:200 |
| **CD19** | SJ25C1 | BUV496 | BD Biosciences | 612938 | 1:200 |
| **CD56** | NCAM16.2 | BUV563 | BD Biosciences | 612928 | 1:100 |
| **CD57** | QA17AO4 | BV510 | Biolegend | 393314 | 1:100 |
| **CD117** | 104D2 | BV711 | Biolegend | 313230 | 1:100 |
| **CD34** | 561 | PE Dazzle | Biolegend | 343534 | 1:100 |
| **CXCR4** | 12G5 | PE-Cy7 | Biolegend | 306514 | 1:100 |
| **NKP80** | REA845 | APC | Miltenyi | 130-112-591 | 1:100 |
| **CD16** | 3G8 | BV650 | Biolegend | 302042 | 1:150 |
| **CD103** | Ber-ACT8 | BV785 | Biolegend | 350230 | 1:150 |
| **CD49A** | TS2/7 | FITC | Biolegend | 328308 | 1:100 |
| **CCR7** | G043H7 | BV421 | Biolegend | 353208 | 1:100 |
| **CD69** | FN50 | PE | Biolegend | 310906 | 1:100 |
| **CD62L** | DREG-56 | AF700 | Biolegend | 304820 | 1:100 |
| **EOMES** | WD1928 | APC | Invitrogen | 50-4877-42 | 1:100 |
| **GRANZYME B** | GB11 | BV421 | Biolegend | 515408 | 1:100 |

**Supplemental Table 1: Antibody panel used for cyclic immunofluorescence of FFPE pediatric tonsil sections**

| ANTIGEN | CLONE | VENDOR | CATALOG # | CONC (UG/ML) |
|---|---|---|---|---|
| **KI67** | B56 | Fluidigm | 3172024B | 5 |
| **CXCL12** | 79018 | RnD | MAB350-SP | 8 |
| **CD3** | Polyclonal | abcam | AB5690 | 1.5 |
| **CD56** | Polyclonal | RnD | AF2408 | 2.5 |
| **CXCR4** | UMB2 | abcam | ab124824 | 2 |
| **CD34** | QBEnd/10 | Novus | NBP2-34713 | 5 |
